# Rheinheimera sp. T2C2 Bacterial Biofilm for Bioremediation of Cobalt (II)

**DOI:** 10.64898/2026.01.21.700925

**Authors:** Ellen W. van Wijngaarden, Miranda P. Brunette, Alexandra G. Goetsch, Ilana L. Brito, David M. Hershey, Meredith N. Silberstein

**Affiliations:** Sibley School of Mechanical and Aerospace Engineering, Cornell University, Ithaca, NY; Department of Material Science and Engineering, Cornell University, Ithaca, NY; Department of Bacteriology, University of Wisconsin-Madison, Madison, WI; Meinig School of Biomedical Engineering, Cornell University, Ithaca, NY; Engineered Living Materials Institute, Cornell University, Ithaca, NY

**Author notes:** Phone: 607/255-5063. Supplementary Information (SI) and supplementary data available at: zenodo.org DOI: 10.5281/zenodo.18225175.

## Abstract

Toxic metals, including cobalt, are often the cause of contamination of rivers and lakes in mining regions. Heavy metal water pollution has been linked to numerous human health problems, prompting the need for environmental remediation. Existing techniques for removing heavy metals from water, such as chemical precipitation and filtration, produce toxic waste, are costly, or require high power consumption for pumping. Biosorption is a potential alternative strategy that is cost-effective and uses readily available and naturally produced biomass and living material to absorb pollutants. Engineering living materials, such as biofilms, which consist of living cells and a secreted polymer matrix, offer potential to integrate toxin sensing, sequestration, and metabolism capabilities of cells to improve pollution remediation strategies. New biofilm producing candidates need to be explored to implement these material capabilities. Previous biosorption studies have primarily used bacterial biofilms from known pathogens and/or generate toxic waste in the form of the absorbent material combined with the heavy metal. Here, we describe a newly isolated bacterium called Rheinheimera sp. T2C2 that forms biofilms with promising biosorption characteristics. T2C2 is a non-pathogenic, aquatic bacterium with low nutrient requirements and high biofilm production. We demonstrate 1) the efficacy of Rheinheimera sp. T2C2 as a biosorbent for cobalt bioremediation; 2) how biosorption is altered by water conditions to establish the efficacy of this strategy in different environments; and 3) how the metal can be released from the biofilm for metal recycling. Our findings will provide a living materials strategy that overcomes existing barriers for bioremediation, and improve the health of ecosystems and humans through heavy metal removal and recycling.

## Introduction

The widespread use of electric vehicles and electronics in daily life has dramatically increased the demand for heavy metal mining. Mining production of cobalt, a key metal for electronics, was over 2.3E5 metric tons in 2023, and above 3E5 metric tons in 2024 [44]. The European Union estimates that only around 32% of cobalt is recycled, driving the high demand for increased mining and with it, heavy metal pollution and negative health effects.[23] Numerous studies demonstrate the toxic effects of heavy metal contamination, including respiratory issues, pancreatic damage, and harm to reproductive organs due to heavy metal ingestion from sources, such as drinking water.[17, 68, 16, 57] The World Health Organization regulates cobalt levels in drinking water to 50 *µ*g/L, although the organization notes that no safe level of cobalt exists.[71] Cobalt concentrations of up to 34.4 g/L have been measured in mine water and concentrations from 0.5 - 2028 *µ*g/L have been routinely measured in surface water in Canada, with values likely to increase further due to mining trends.[72]

The increased demand for cobalt mining and the negative health effects due to heavy metal pollution necessitate remediation strategies, however many of these techniques have drawbacks. Chemical precipitation is simple and cost-effective, but produces toxic waste and only works in a narrow pH range.[16, 36] Ion exchange successfully recovers metal at low concentrations in the parts per billion range.[24, 16, 36, 1] However, ion exchange is an expensive method for high volumes of effluent, particularly at high metal concentrations that quickly saturate metal binding sites. Physical methods, such as filtration, are costly, require regular anti-fouling maintenance, or need high power consumption for pumping.[51, 16] Ultimately, the drawbacks of these chemical and physical strategies for remediation hinder implementation and widespread use.

Bioremediation techniques have gained interest as a means to overcome some of the disadvantages of current remediation methods. Biosorption presents a cost-effective strategy with minimal toxic waste due to the use of natural, readily available, and often biodegradable material. Bioderived and engineered living materials (ELMs) present exciting possibilities for enhancing biosorption, to include toxin sensing[69, 64, 38, 3] and sequestration[16, 5]. Current bioremediation techniques use biomass from an array of sources, including plant matter, mushrooms, algae, or biofilms.[54, 40, 60, 46, 27, 14, 52, 13, 45, 11, 19] While most of these examples are primarily academic demonstrations, algae has been deployed commercially for wastewater remediation technologies.[58] Biosorption using algae packed in columns has proven to be effective, however this method still requires the disposal of algae or biosorbent with absorbed metal and synthetic biology tools for working with algae remain limited. Therefore, routes to improve the system from a biological engineering end are limited.[54] Bacterial biofilms have also proven to be quite effective for metal uptake.[7] These biofilms are a combination of live cells and self-produced extracellular polymeric matrix, typically rich in high molecular weight (MW) polysaccharides, proteins, lipids, and DNA.[18] Synthetic biology along with tuning environmental factors, such as growth conditions, offer multiple strategies to improve biofilm yield, metal uptake, and biosafety, thereby supporting system level engineering.

Widespread implementation of bacterial biofilm-based ELMs for metal recovery requires wild type, native, non-pathogenic bacteria that produce high amounts of biofilm, and achieve effective metal removal. Previous work has focused on bacteria that meet only some of these requirements, including prolific biofilm producers, such as *Pseudomonas aeruginosa*.[16, 62] While *P. aeruginosa* is pathogenic, which poses a barrier to widespread use, it has been proven to be effective for heavy metal uptake.[16, 62] Previous work has demonstrated that bacteria of the genus *Geobacter* can take up cobalt and down-cycle it. While this does not allow for metal recovery to decrease the demand for mining, this strategy might be preferable in environments where recovery is not possible or contamination is exceptionally widespread.[19] The use of wild-type, non-pathogenic bacteria, such as soil strains, for heavy metal uptake, holds great potential for widespread use and has been demonstrated in a contained environment with industrial level waste concentrations. This direction is promising but requires additional study in varied environments and lower pollution level concentrations.[22] We have taken inspiration from this previous work to explore the properties of an aquatic bacterium, ideal for water remediation.

An ELM for bioremediation, specifically for cobalt pollution, requires a nonpathogenic, aquatic organism with stable growth and consistent rheological properties across temperature and pH ranges for water environments, such as lake water where cobalt pollution is high.[72] Additionally, high biofilm production under low nutrient conditions is needed to maintain the material. Differences in the composition and structure of the biofilm matrix may lead to fundamental differences in bioremediation performance between organisms. For example, the abundance of hydroxyl and carboxyl functional groups, a compositional characteristic, would be ideal for metal adsorption. Such an organism would greatly facilitate further design and optimization for metal uptake based on the environmental conditions and exposure time. Here, we describe a newly isolated[25] bacterium called *Rheinheimera sp.* T2C2 that produces a biofilms with promising properties for bioremediation of cobalt. We demonstrate 1) the efficacy of *Rheinheimera sp.* T2C2 biofilm as a living material for cobalt bioremediation; 2) how biosorption is altered by water conditions and exposure time to establish the efficacy of this strategy in different environments; and 3) an approach for releasing metal from the biofilm for metal recycling.

## Results and discussion

### *Rheinheimera sp.* T2C2 as an Ideal Candidate for Heavy Metal Bioremediation

Ideal bacteria for use in ELM development would be non-pathogenic, wild-type, aquatic, and able to survive with low nutrients to ensure viability under varied conditions (e.g. temperature, pH, nutrients) of use. Although bacterial biofilms have been shown to have potential for bioremediation of pollutants, not all bacteria are ideal candidates. *Rheinheimera* is not known to be pathogenic and has even been found to inhibit the growth of pathogenic bacteria.[8, 10] *Rheinheimera sp.* T2C2, first isolated from a water sample from Lake Mendota (Madison, WI), demonstrates the qualities of being an ideal candidate for bioremediation.[25] The genus *Rheinheimera* includes various aquatic bacteria found in both saltwater and freshwater, as well as soil all over the world.[67, 9, 49, 56, 63] These bacteria have low nutrient requirements, and therefore are typically grown in the lab setting on peptone yeast extract (PYE) media.[56, 26] Previous studies have mapped out the phylogeny of various strains and identified a range of ideal growth conditions depending on the strain.[56, 67, 70] These include strains that have been shown to grow well between 10*^◦^*C and 40*^◦^*C and at a pH between 7 and 8, conditions typical of lake water environments.[9, 56] Reported lake water conditions can range from approximately 9*^◦^*C to 30*^◦^*C and pH values from 6 to 8 according to the National Oceanic and Atmospheric Administration.[59, 53] While biofilm could be grown in ideal conditions before transfer to the environment, cell viability and continuous production of biofilm or intracellular transport of metal could be advantageous for metal removal and continuous remediation. We show that *Rheinheimera sp.* T2C2 grows well at both 20*^◦^*C and 30*^◦^*C with minimal growth at 7*^◦^*C in PYE media (Figure 1a, Figure S1). Extended growth for multiple days slightly improves biofilm yield but results in a lower optical density value, likely due to autolysis (Figure S1a,b).

**Figure 1:**
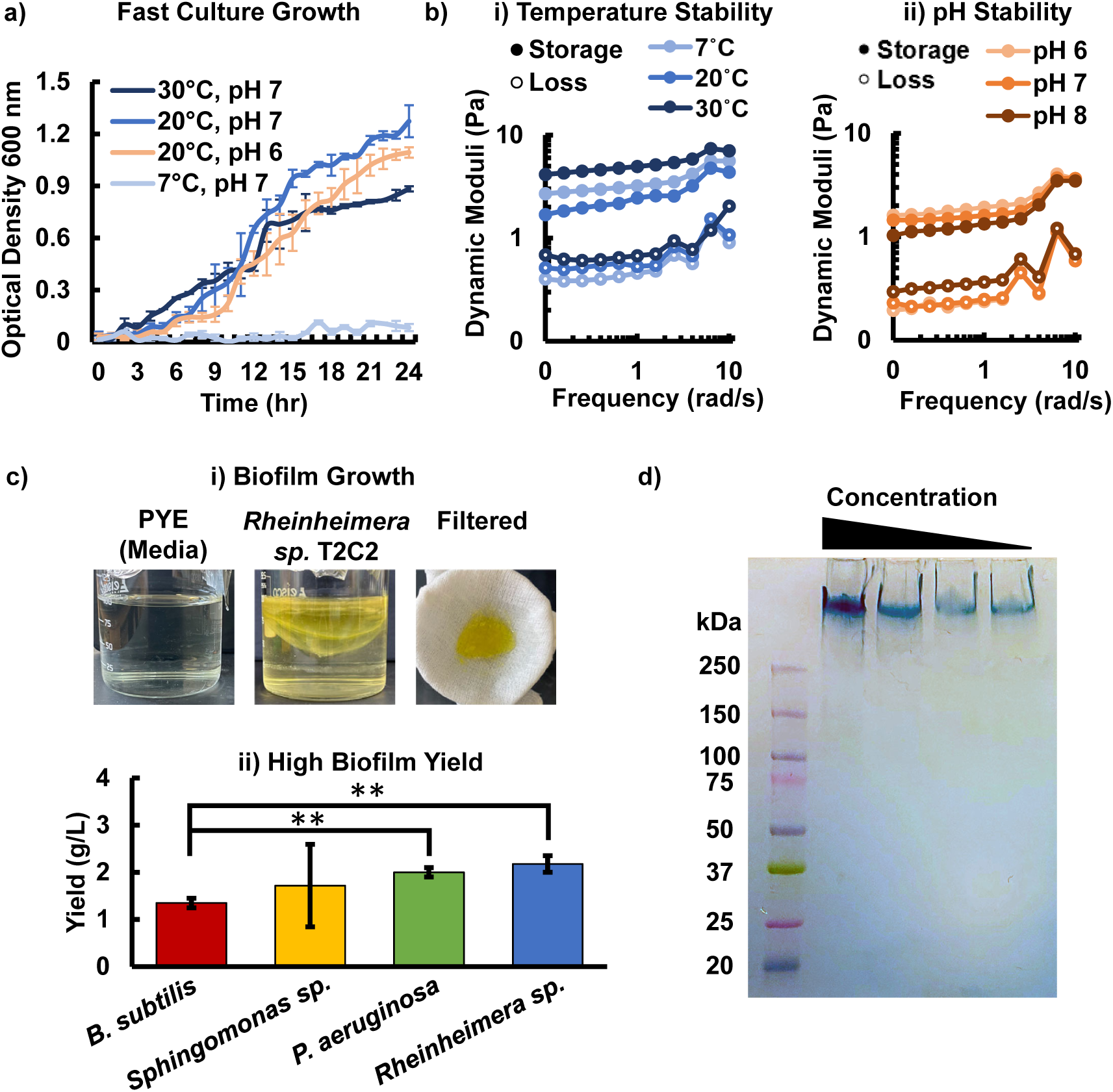
*Rheinheimera sp.* T2C2 biofilm is an ideal biosorbent due to its a) fast growth in varied environmental conditions with low nutrient requirements and b) stable biofilm rheological properties under different environmental temperatures and pH values. c) The formation of insoluble biofilm with a d) high yield compared to other model biofilm producing bacteria, such as *B. subtilis*[21], *Sphingomonas sp.*[65], and *P. aeurinosa*[43], would facilitate the use of *Rheinheimera sp.* T2C2 for bioremediation. Error bars show standard deviation, n=3, **p-value *<* 0.01 (two-tailed student t-test). e) Alcian blue staining indicates that the biofilm is acidic providing further evidence for potential for metal binding.

The growth and environmental conditions may also alter the biofilm properties. Rheology was conducted to check the stability of the biofilm at varied environmental conditions. Material stiffness may play a key role in metal diffusivity and consistent properties are desirable for implementation of a living material. Figures 1b, and Figure S2a, b show a stable gel behavior for all lake water relevant temperatures and pH values with minimal change in the dynamic moduli between conditions. Our results also show growth for pH values of 6 and 7 (Figure 1a). These results indicate that the living material would have consistent properties if applied in various lake water-relevant conditions.

The high biofilm yield of *Rheinheimera sp.* T2C2, compared to other model organisms is also ideal for bioremediation. T2C2 produces a higher amount of biofilm than *Sphingomonas sp.* LM7, a member of the *Sphingomonas* genus that are often harvested for use as rheological modifiers for use in food, oil and gas, and cosmetics, using the same PYE growth media.[65] Further, our results show that *Rheinheimera sp.* T2C2 produces more insoluble biosorbent material (1c) than other high biofilm producers, such as *B. subtilis* (Figure 1d).[43, 21] T2C2 biofilm yield was similar to model biofilm producer *Pseudomonas aeruginosa* 215 but without the risk of pathogenicity that *P. aeruginosa* is known for. Note that the examples for *B. subtilis* and v*P. aeruginosa* are in their respective preferred media and growth conditions.

The composition and structure of the biofilm produced, indicates that this material might be particularly well-suited for metal uptake for bioremediation applications. The amount of metal uptake would depend on environmental conditions and factors related to the metal, such as oxidation state and electronegativity, in addition to biofilm composition.[28] The biofilm is comprised primarily of polysaccharides with additional protein, RNA, DNA, and lipids (Figure S3a). The biofilm is approximately 5 % protein by dry weight (Figure S3a) and is mainly composed of high-molecular weight polysaccharides. The purified polysaccharide molecular weight was approximately 11.7 MDa with a surface charge or zeta potential value of -44.5 ± 2.4 mV. The low zeta potential value, below -30 mV, indicates that the polysaccharide molecules will repel one another rather than aggregate in solution.[37, 42, 65] This is an ideal characteristic for a bioremediation to ensure biofilm mixing with a desired heavy metal or toxin rather than flocculating out of solution. The acidic nature of the extracellular polymers, as shown in Figure 1e may also help with metal binding as acidic polymers can donate electrons to metal ions to form stable bonds.[15, 29] The blue color of the Alcian blue staining indicates acidic functional groups, which were further explored using FTIR (Figure S3b). Key functional groups identified, such as amides, carbo-acids, and alcohols, may facilitate binding to metal ions (Figure S3b). T2C2 shows strong stretching for amide I for C=O at 1680-1630 cm^-1^, NH at 1580-1515 cm^-1^, alcohol groups for C-O at 1075-1000 cm^-1^, and C=O from 1670-1650 indicative of internal hydrogen bonding. The presence of carboxyl, amine, and hydroxyls indicate potential for metal binding and effectiveness for bioremediation. The combined compositional evidence points towards the potential of T2C2 biofilm for metal uptake.

### Biosorption of Cobalt using *Rheinheimera sp.* T2C2

Cobalt biosorption was tested to compare T2C2’s performance to other biosorbents. T2C2 biofilm pellets were exposed to water containing varied concentrations of cobalt chloride salt, within an environmentally relevant range, in a transwell plate. The transwell membrane size of 0.45 µm prevented any cells from migrating into the water below the transwell but the metal ions were free to diffuse into the biofilm in the upper part of the transwell (Figure 2a inset). A colorimetric reaction with nitroso-R salt was used to measure the change in cobalt concentration in the free solution following biosorption (Figure S4). Increasing concentrations of cobalt resulted in a lower percentage metal removal as metal binding capacity of the biofilm was saturated at higher concentrations (Figure 2a). However, the overall amount of metal removed was still higher due to the increase in initial solution concentration.

**Figure 2:**
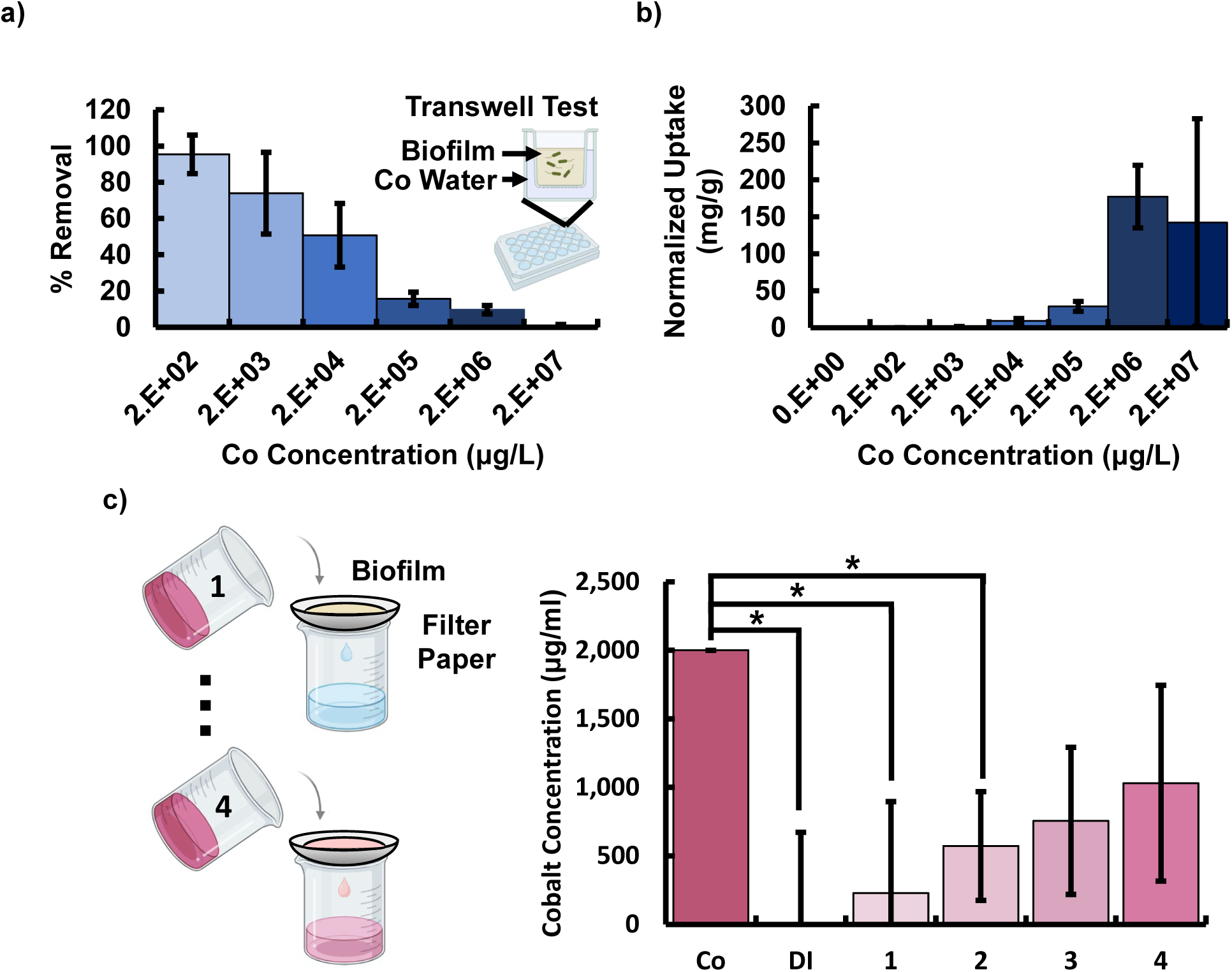
*Rheinheimera sp.* T2C2 demonstrated effective bioremediative properties showing effective a) removal of cobalt from water and b) metal uptake normalized by biofilm mass for a range of environmentally relevant metal concentrations spanning 200 *µ*g/L to 20 g/L. c) Metal could also be removed via filtration using biofilm pellets to achieve comparable uptake to the transwell experiments while decreasing exposure time. Cobalt contaminated water was poured through a coffee filter containing the biofilm pellet in 10 ml increments and the volumes of filtered water were collected individually for cobalt measurement. The initial cobalt concentration was 2E+03 *µ*g/L. Tests were performed in triplicate, error bars show standard deviation.

The normalized uptake of the biofilm (Figure 2b) was determined to add insight into the biofilms effectiveness at different cobalt concentrations and to facilitate comparisons with other biofilm producers. Normalized uptake, sometimes referred to as uptake capacity (*Q*), was calculated by multiplying the difference in the initial (*C_initial_*) and final metal concentrations (*C_final_*) by the volume of cobalt contaminated water (*V*) and dividing by the biofilm mass (*M*):

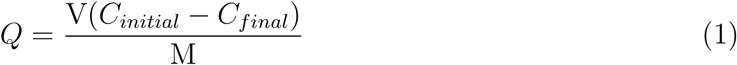

A peak value of uptake was obtained for 2E+06 µg/L of cobalt, with metal uptake likely saturating above this concentration. At 200 µg/L, the Q-value for T2C2 was 175 µg/g (95 % removal) compared to *P. aeruginosa* which took up 1.19 µg/g (25 % removal) under similar initial metal and water conditions (pH 7, room temperature, and 24 hours of exposure).[16] Uptake for another high performing biosorbent reported in literature, seaweed derived from *S. wightii*, was 16 mg/g for an initial cobalt concentration of 100 mg/L (16 % removal) and 12 hour exposure. T2C2 showed comparable metal uptake of 13 mg/g but at a lower initial cobalt concentration of 20 mg/L (65 % removal) for the same exposure time, indicating better bioremediation capacity.[61]

Uptake can also be achieved with short exposure by running contaminated water through a coffee filter containing a biofilm pellet, presenting a possible alternative strategy for gravity driven filtration that does not require long exposure periods. We demonstrated this option by placing the biofilm pellet on the coffee filter while 10 ml volumes of cobalt contaminated water were incrementally filtered and collected. This technique yielded significant metal uptake for the initial samples collected. Metal percent removal was comparable to the transwell tests due to the high amount of biofilm used. As expected, the percent of metal removal declined for repeated exposure to cobalt contaminated water, indicating a gradual saturation of uptake sites (Figure 2c). Debris from the coarse coffee filter likely contributes to the high error. The first volume filtered allowed only 229 *µ*g/L through the filter or 11% metal remaining, while the fourth volume had a filtered concentration of 1030 *µ*g/L or 52% metal remaining. This demonstrates how bioremediation could be scaled to increase filtrate volume and reduce time given the high biofilm yield and low cost of production.

It is important to note that high cobalt concentrations can be cytotoxic to bacteria, hindering material production in a living material, which is essential for continued metal sorption. We found that cells were capable of growing biofilm in the presence of low concentrations of cobalt. The biofilm had consistent water content (Figure S5a), yield (Figure S5b) and rheological properties (Figures S5c,d) at cobalt concentrations up to 2E+03 *µ*g/L. Without initial biofilm matrix to protect cells, 2E+04 *µ*g/L was too high of a cobalt concentration (Figure S5a,b) for cell growth. However, already grown cells, within a mature biofilm matrix, could withstand higher cobalt concentrations (Figure S6a). We measured cell viability after exposure in the transwell plate for cells with a mature matrix prior to cobalt exposure. Cobalt had minimal impact on cell viability up to a concentration of 2E+04 µg/L (Figure S6a), at which point there was a dramatic drop off with increased concentration. FTIR-ATR results indicated that the biofilm matrix is not just important for cell viability but also for metal uptake, clearly visible at high concentrations (Figure S6b). Results show changes in the peaks, particularly in the protein region between 1580-1515 cm^-1^, indicating metal binding to these groups. Since our results indicate that cells without a mature matrix are susceptible to cobalt exposure, growing the biofilm in advance under ideal conditions before exposure might be a more effective bioremediation strategy than growing biofilm in contaminated water, to better protect cells and promote metal uptake.

### The Effect of Varied Water Temperature and pH on Biofilm Biosorption of Cobalt

Temperature and pH must also be considered when designing a living material for bioremediation. Living organisms may respond adversely to temperature or pH extremes, with environmental conditions likely influencing cell viability and perhaps also metal uptake. While temperature and pH are often beyond human control, exposure time could be potentially modulated to achieve effective uptake. We tested a range of environmental conditions to determine how temperature and pH affect metal uptake. The role of exposure time was then examined in addition to these environmental factors.

Varied water temperature conditions have been shown to affect uptake performance of biofilms.[16] We therefore examined uptake at three temperatures, 7*^◦^*C, 20*^◦^*C, and 30*^◦^*C, chosen based on the range of temperatures observed in lake water.[41, 33] We first grew biofilm under consistent ideal conditions before exposing to the three temperatures of interest. We observed metal uptake at all conditions tested (Figure 3). For 6 hours exposure time, our results show the highest uptake for 20*^◦^*C. However, all temperatures eventually reached similar metal percent removal by 24 hours (Figure 3b, Figure S7a). At the 24 hour time point, temperature did not alter the percent removal (Figure 3, Figure S7a). Lower metal percent removal was observed for higher concentrations of cobalt, indicating a gradual saturation of cobalt binding sites (Figure 3a). We observed the fastest cobalt removal at 20*^◦^*C with metal concentrations already statistically significantly altered by 6 hours for 2E+03 *µ*g/L cobalt in wells with biofilm compared to wells without biofilm at 20*^◦^*C (Figure 3b). All concentrations of cobalt showed statistically significant metal uptake by 12 hours at 20*^◦^*C. In contrast, samples at 7*^◦^*C showed statistically significant uptake by 12 hrs at a concentration of 2E+03 *µ*g/L and samples at 30*^◦^*C did not show any statistically significant uptake until 24 hrs (Figure 3b). Changes in temperature may influence several factors including metal solubility, bacterial cell wall configuration, ionization energy of metal-biomass complexes, diffusivity, and reaction equilibrium. Metal coordination complexes can be formed endothermically or exothermically. For example, metal cations and carboxylate ligands form a complex endothermically whereas, amide ligand complexes form exothermically.[34, 30] This might obscure clear trends for metal uptake and contribute to high variation in data. Previous studies have indicated that an increase in temperature can decrease the capacity for physical adsorption, aligning with our observations that 30*^◦^*C results in lower metal uptake than 20*^◦^*C initially.[48, 6] However, this does not appear to have an effect at longer exposure times. Viability was highest for 20*^◦^*C for all concentrations and time points (Figure S7b). The number of colony forming units per milliliter (CFU/ml) gradually decreased with increasing exposure time and increasing cobalt concentration, indicating the need for investigating the role of active cell uptake compared to cell surface or biofilm sorption.

**Figure 3:**
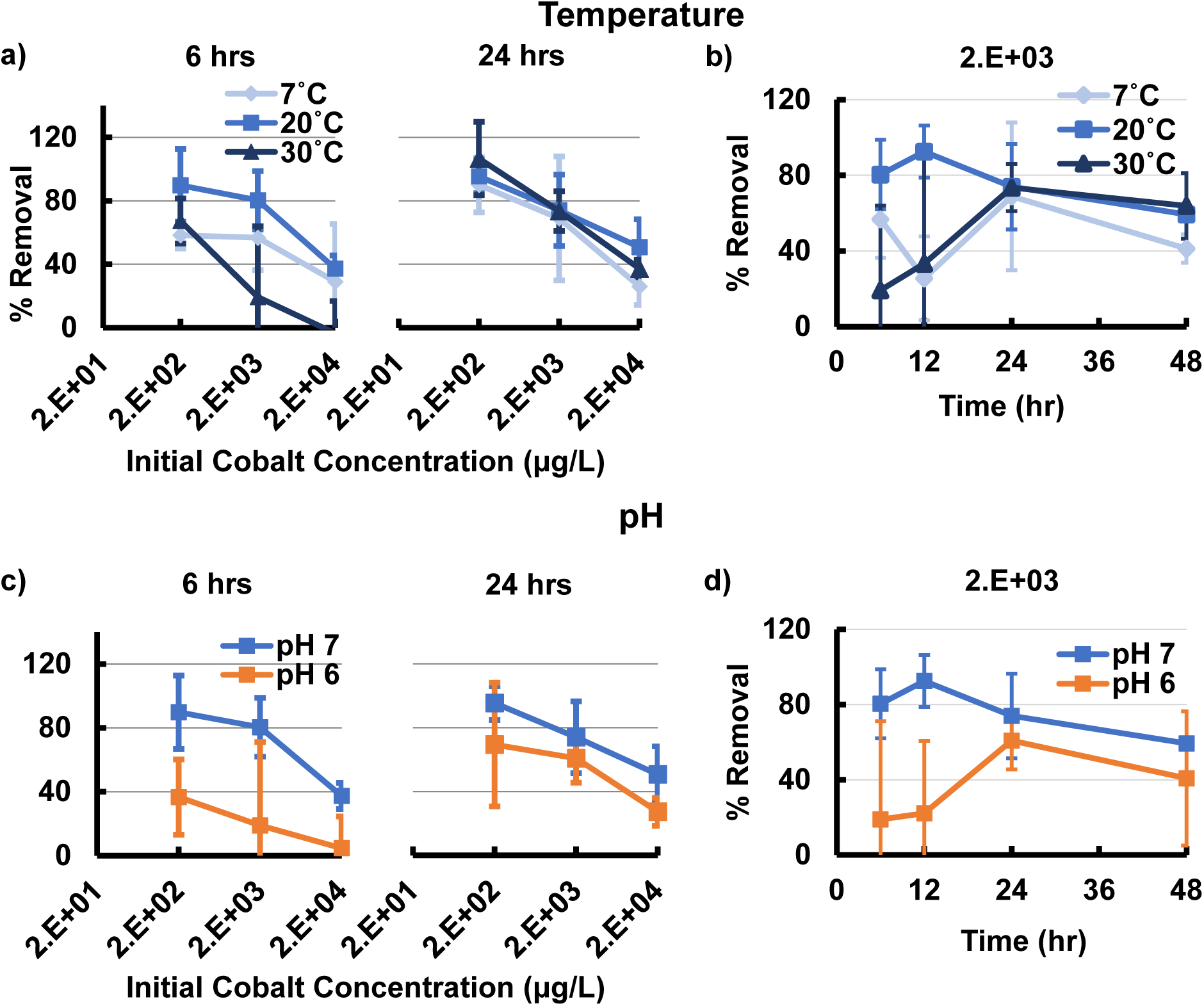
a) The effect of varying water temperature for exposure times of 6 hrs and 24 hours. b) Varying water temperature showed varied percent cobalt removal peak uptake times. The effects of temperature lessen for longer exposure times. c) The effect of varying pH for exposure times of 6 hrs and 24 hours, on the percent cobalt removal showing a decreasing trend with metal concentration. d) The effect of varying water pH and exposure time on percent cobalt removal showed differing peak uptake times depending on pH. Error bars show standard deviations (n=3).

Varied water pH also affected uptake performance of the *Rheinheimera sp.* T2C2 biofilm grown at standard conditions. The pH of the surrounding water may alter the nature of binding sites and the solubility of metals. Biofilms may contain both weakly acidic and weakly basic groups depending on composition.[55] A decrease in pH increases the H^+^ ions in the solution, leading to a positive charge on the cell and biofilm surfaces, which could inhibit attraction between metal and biomass.[7, 48] In contrast, an increase in pH has been shown to increase biofilm metal normalized uptake due to the net negative electrostatic surface charge, matching our observations.[48] For cobalt (II), the optimal pH, according to literature, is between 4 and 7, as a pH of approximately 8 and above results in cobalt removal by hydroxide precipitation, creating high volumes of sludge that is difficult to dispose of.[48, 35] This process interferes with cobalt removal by adsorption.[35] We explored the effect of pH on metal uptake of our living material by testing pH values of 6 and 7 to match the pH of typical lake water.[41] Previous studies have explored broad pH ranges to establish trends in metal binding and indicated that a pH close to neutral is ideal.[16, 48, 54]

We observed higher uptake for pH of 7 in comparison with pH 6, except for at a metal concentration of 200 *µ*g/L and an exposure time of 48 hours (Figure S8a). A downward trend in percent metal removal with increased concentration of cobalt was observed for both pH values (Figure 3c). The time for percent removal to equilibrate was higher for a pH of 6 compared to a pH of 7, particularly at lower concentrations (Figure 3d). The drop in metal removal for 48 hours compared to shorter exposure times may be due to the degradation of biofilm over time.[48] Viability was higher for a pH of 6 compared to a pH of 7 for all concentrations and exposure times (Figure S8b, S9b) indicating that more cells does not necessarily lead to higher metal removal and that pH can alter multiple factors that are important for bioremediation.

### Biofilm Mode of Cobalt Uptake

Establishing the mode of uptake may identify approaches to improve cobalt removal as well as possible cobalt recovery strategies. Metal uptake may occur due to intracellular accumulation, cell surface sorption, and extracellular accumulation. Extracellular vs. Intracellular uptake is challenging to study as transport of heavy metals into the cell often leads to toxicity and cell death. Cells have mechanisms for uptake of essential ions such as sodium and potassium, that may also lead to uptake of heavy metals.[16] Cell surface and extracellular adsorption is likely due to binding with functional groups present on the cell membrane and secreted biofilm, including hydroxyl, carboxyl, and amino groups (Figure S6b).[68, 16] These functional groups contribute to the negative surface charge of the T2C2 biofilm matrix, with a zeta potential (-44.5 ± 2.4 mV) comparable to other bacterial polysaccharides, such as *Sphingomonas sp.* LM7 with a surface charge of -31.8 ± 2.7 mV and gellan gum with a value of -29.1 ± 3.0 mV.[2, 65]. The negative surface charge may assist with uptake of positively charged cobalt ions. The nitrogen atom from amine groups and the oxygen atom from hydroxyl and carboxyl groups can bond with the cobalt metal ions.[68, 16, 47] The lower electronegativity of nitrogen, compared to oxygen, leads to a higher bonding efficiency.[47] Different cell surface components may also contribute to uptake reactions via the various functional groups. Lipopolysaccharide (LPS) plays a role primarily in carboxyl and hydroxyl reactions. Protein is also involved in most cell surface adsorption events as reported for *Pseudomonas aeruginosa*.[34] Additionally, phospholipids have also been identified as a key component for hydroxyl-metal binding, demonstrating the vast array of binding mechanisms.[32, 12, 20]

We conducted separate metal uptake tests with cells and biofilm to assess the role of active cellular uptake and determine whether extracellular matrix or cells contributed more to metal removal for *Rheinheimera sp.* T2C2. The biofilm was centrifuged to separate the matrix from the cells before conducting metal removal experiments. Results indicate that the matrix portion had statistically significantly more cobalt uptake than the cell component (Figure 4a). Note that centrifugation did not completely separate the matrix and cell components as some colonies still grew when matrix was plated (Figure S10a). Separating the matrix and the cells made the cells much more susceptible to cobalt exposure. Cobalt exposure killed all separated cells, which were not protected by a biofilm matrix, and also resulted in significant cell death to cells still embedded in the matrix (Figure S10a). This could be due to the combined stress of centrifuging the biofilm at high speed (7830 rpm) and the metal treatment. Note that a higher speed was used here to actually separate cells from film, not merely collect biofilm from culture solution. The overall number of cells in the matrix and cell components added to approximately the same as the whole biofilm, indicating that centrifuging alone did not significantly affect viability. Separated cells were treated with lysozyme to lyse cells and eliminate active intracellular uptake but preserve cell surface sorption to determine how physiological state, living versus dead, altered cobalt uptake. Note that lysozyme was ineffective for lysing cells without prior centrifugation and separation (Figure S10b) as the extracellular matrix protected cells embedded within pores,[66] resulting in no change in the percent cobalt removed (Figures S10b,c). Separating the matrix component did not significantly alter the FTIR spectra compared to the intact biofilm (Figure S10d). Together, these tests indicate that the intracellular uptake does not play a large role in cobalt removal, supporting that uptake is primarily due to functional groups on the extracellular matrix or cell surface. The cobalt uptake due to the matrix portion was not statistically significantly different from the cobalt removal measured for the biofilm as a whole (Figure 4a).

**Figure 4:**
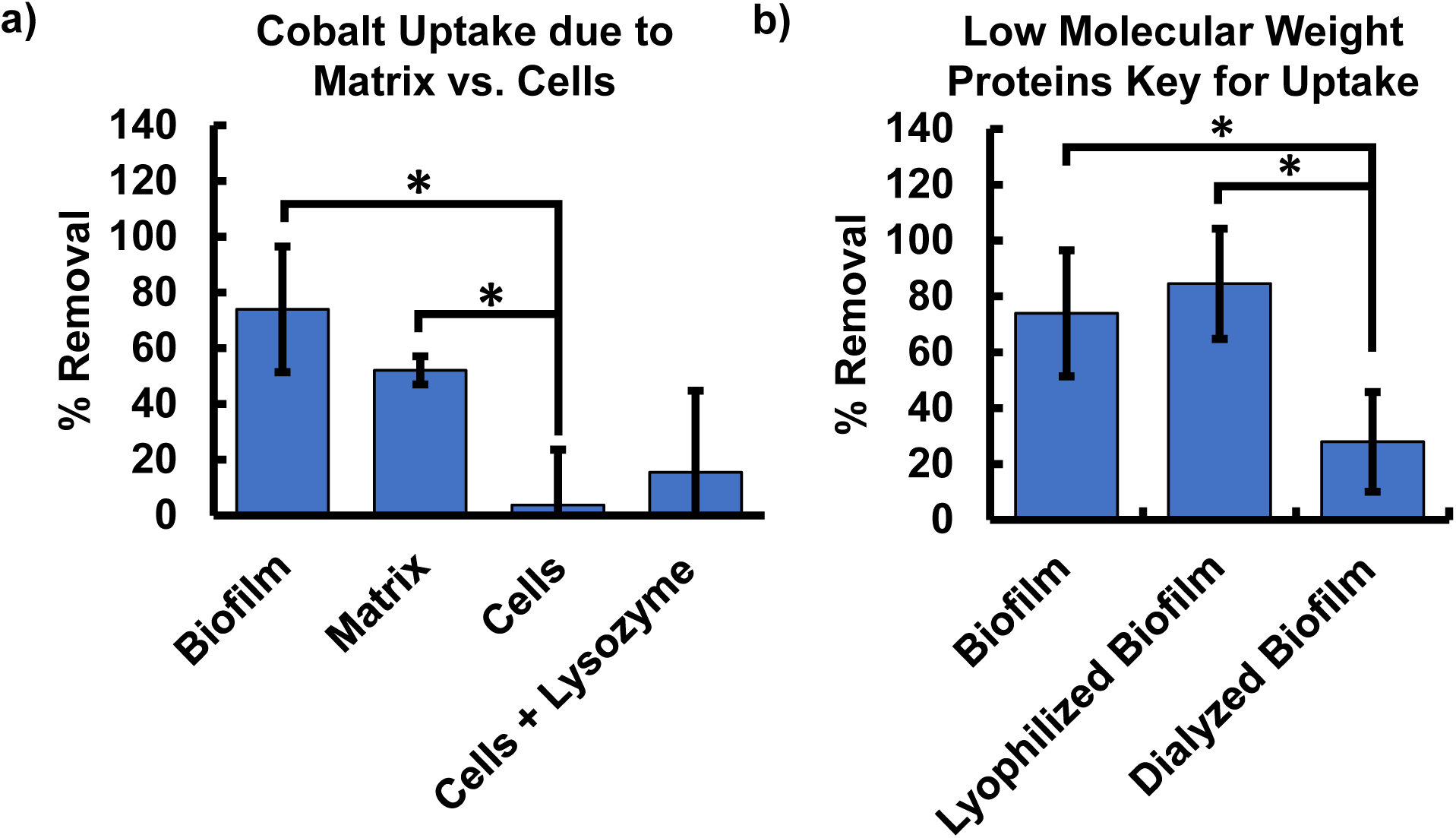
a) The mode of cobalt uptake (intracellular vs. extracellular) was explored by separating and lysing cells from the biofilm before measuring metal uptake, demonstrating that uptake was primarily due to the matrix component. b) Lyophilization and 100 kDa dialysis post lyophilization were used to kill all cells and eliminate low molecular weight proteins, respectively, to determine the component (polysaccharide or proteins) of the matrix that contributed most to cobalt uptake. Plots report mean values with error bars showing standard deviations (n=3).

Based on the prior determination that the extracellular matrix was more critical than the active cells for uptake, we set out to determine what component of the extracellular matrix was most important. Specifically, to determine whether matrix polysaccharide or proteins were the primary components for cobalt uptake. To isolate the effect of the matrix metal uptake without living cells, biofilm was lyophilized, resulting in complete cell death without the addition of any chemicals, such as lysozyme. Dialysis was used, following lyophilization, to eliminate proteins under 100 kDa, to investigate which component of the matrix contributed most to uptake, low molecular weight proteins or the high molecular weight polysaccharide. No significant difference was observed between biofilm cobalt removal that was lyophilized and initial biofilm, indicating that lyophilization did not alter the biofilm structure or composition in a way that would affect cobalt uptake. In contrast, 100 kDa dialysis for 24 hours did result in a statistically significant decrease in cobalt removal, indicating that low molecular weight proteins play a significant role in cobalt binding (Figure 4b, Figure S3). Note that proteins may play a direct role in binding metals or have an indirect effect via altering polysaccharide conformation to affect metal binding.[4] This experiment identifies proteins as a key biofilm component for cobalt uptake.

### Cobalt Recovery via Biofilm Protein Degradation

Due to the key role of proteins in the cobalt uptake of *Rheinheimera sp.* T2C2, breaking down proteins presents a possible strategy for cobalt recovery for recycling purposes. Biofilms were treated with Proteinase K and guanidinium iodide to explore targeting proteins as a strategy for cobalt recovery. These chemicals primarily target the protein component of the biofilm and are more cost-effective than the use of enzymes to break down the polysaccharide component of the biofilm, a strategy that is also limited by specificity and understanding of the linkages and bonds comprising the polysaccharide structure. Proteinase K is a serine protease used to digest proteins and remove enzymes that break down nucleic acids. It is known as a broad spectrum protease and is capable of breaking down a variety of proteins by bonding to the carboxylic side of both aromatic and aliphatic amino acids to digest and inactivate proteins and enzymes. Guanidinium iodide and other guanidinium salts have been identified as effective protein denaturants.[31] Literature indicates that guanidinium salts primarily target proteins but may also degrade or alter polysaccharide and other biopolymer structures.[31] We hypothesized that these methods of digesting and disrupting the protein structure could be used for reversing metal binding for possible recovery.

Our results show significant changes in the biofilm structure and cobalt binding when exposed to Proteinase K or guanidinium iodide. Treatment with either chemical significantly reduces the dynamic moduli of the biofilm gel, indicating structural changes to the biofilm (Figure 5a, Figure S11a). Subsequent chemical treatment, for biofilm that had uptaken cobalt, showed that cobalt could be recovered from the biofilms (Figure 5b). Guanidinium treatment resulted in a statistically significant amount of cobalt recovery into the free solution. Exposure to guanidinium also resulted in 100% cell death, whereas, the majority of cells survived Proteinase K treatment (Figure S11b). Proteinase K also resulted in the release of metal into the free solution, however, the change was not statistically significant for our initial transwell experiments (Figure S11c). Guanidinium treatment resulted in 100% cell death, whereas, the majority of cells survived Proteinase K treatment (Figure S11b). Due to the toxicity of guanidinium, Proteinase K may pose a preferable metal recovery strategy. Proteinase K efficacy is known to improve with temperature, with optimal performance between 20*^◦^*C and 60*^◦^*C.[50] Therefore, conditions could be further altered to optimize chemical treatment for maximum metal recovery.

**Figure 5:**
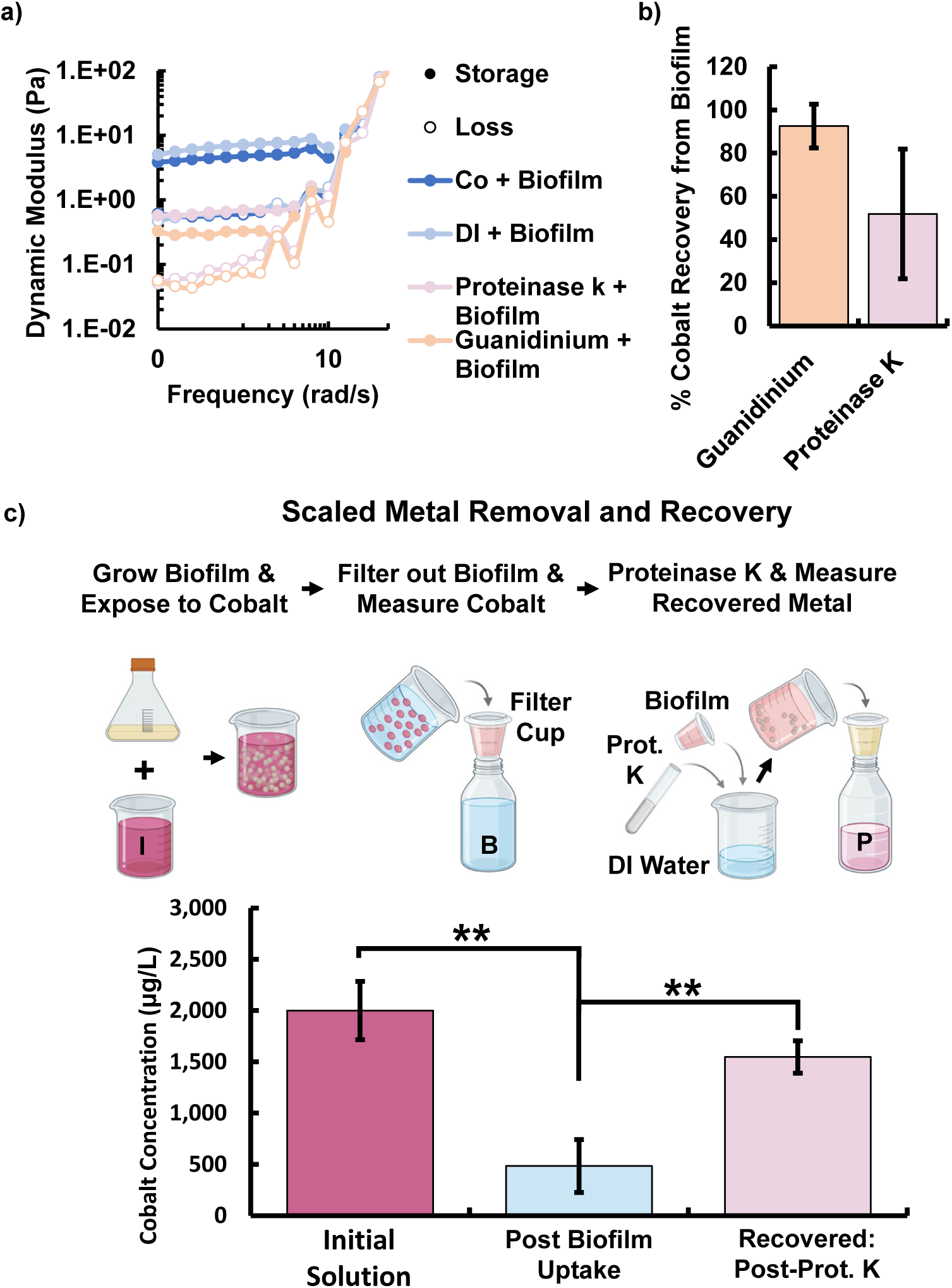
Biofilm was degraded for the purpose of metal recovery using Proteinase K (0.2% w/v) and a protein denaturant, guanidinium iodide (10% w/v) for 6 hours respectively. a) Both chemicals successfully degraded the biofilm, indicated by the decrease in biofilm dynamic moduli, and b) resulted in return of cobalt into the solution. c) Schematic of the cobalt uptake and recovery process performed for liter scale water volumes. The biofilm was grown (2 g dry weight), harvested, and exposed the biofilm to 1 L of cobalt contaminated water at the initial metal concentration. The biofilm was then separated from the free solution to determine the amount of cobalt that was taken up. Finally, the biofilm was treated with Proteinase K in a water bath, filtered out, and the recovered cobalt in the water was measured. The cobalt measurements for these three key steps are shown. All graphs show mean and error bars show standard deviations (n=3).

A scaled cobalt uptake and recovery test was then performed to test the potential for larger volume metal removal and recycling. A liter of cobalt contaminated water was exposed to biofilm at a concentration of 2 mg/ml. After 24 hours of exposure, a sample of the biofilm solution, approximately 100 ml, was extracted and filtered to remove the biofilm. The amount of cobalt removed from the water was measured to infer the amount of cobalt that had been taken up. The biofilm containing cobalt was then treated with Proteinase K to release the metal for measurement of cobalt recovery. The test demonstrated statistically significant uptake and recovery of cobalt using Proteinase K and also showed the potential for liter scale bioremediation (Figure 5).

### Conclusion

Biosorption of pollutants and toxins, particularly heavy metals, offers a promising bioremediation strategy that could reduce reliance on conventional electrochemical and filtration-based methods, which can be energy-intensive, costly, and generate substantial chemical waste. The design of living materials has the potential to elevate toxin remediation to include sensing, sequestration, and metabolism of targeted species. However, existing biosorbents often rely on pathogenic bacteria to achieve sufficient biofilm production, and few systems enable efficient metal recovery to support recycling efforts and minimize toxic biomass disposal. In this work, we investigated *Rheinheimera sp.* T2C2’s performance for bioremediation applications relative to current biomass remediation strategies. The non-pathogenic, wild-type, and aquatic characteristics of T2C2 render the bacterium well suited for biosorption applications. T2C2 exhibits high growth across a range of environmental conditions, possesses matrix functional groups associated with high-affinity metal binding, and achieves superior biofilm yield relative to commonly studied biosorbents. Our results show that T2C2 biofilm demonstrates significantly enhanced cobalt uptake under comparable conditions compared to biosorbents such as *P. aeruginosa* biofilm and seaweed, underscoring its potential as an effective and scalable bioremediant. Moreover, we introduce two recovery strategies that enable cobalt release and collection, providing a preliminary path toward integrated biosorption and recovery for metal recycling. Together, these findings position *Rheinheimera sp.* T2C2 as a compelling new candidate for living material-based cobalt recovery for bioremediation. Continued development of ELMs through leveraging such biofilm-forming organisms will be essential to advance sustainable pollution mitigation and improve environmental well being and human health.

## Experimental

### Rheinheimera sp. T2C2 Isolation

A volume of 1 mL of water from Lake Mendota in Madison, WI was inoculated into 10% strength PYE medium and allowed to incubate at room temperature without shaking. After 2 days, biomass was extracted from the air-liquid interface of the culture, homogenized, serially diluted and plated on PYE agar. A single, mucoid colony was purified by repeated re-streaking on PYE agar. The 16S rDNA gene was amplified using PCR and sequenced using the Sanger method to identify the isolate as belonging to the Rheinheimera clade. A large collection of Rheinheimera strains isolated from Lake Mendota will be described elsewhere.[25]

### Bacterial Culturing

All reagents were sourced from Sigma Aldrich (St. Louis, MO) unless otherwise noted. A 10 mL Peptone Yeast Extract media (PYE: 0.2% w/v peptone w/v yeast extract) with 3% sucrose starter culture was innoculated with LM7 cells from a frozen glycerol stock and grown at 30*^◦^*C overnight on a shaker at 200 rpm. A 10 mL of starter culture was added to 1 L of PYE and 3% w/v sucrose solution. The culture was grown until approximately the end of day 4 at 18*^◦^*C with shaking at 200 rpm. All cultures were grown until an optical density at 600 nm of 0.5 was reached.

### Yield Measurements and Optical Density

Optical density of the cultures at 600 nm was measured using a plate reader (Spectroquant, Darmstadt, Germany). Yield of the material was determined by lyophilizing three 10 ml volumes of culture with a known optical density. Samples were left in the lyophilizer (FreeZone 2.5 L Benchtop Freeze dryer, Labconco, Kansas City, MO) for 24 hours until they were fully dry, before they were weighed to determine yield per volume of culture.

### Metal Uptake Tests

After 4 days of growth following the bacterial culturing procedure described above, the culture optical density was adjusted to 0.5 by adding additional PYE media as needed. The culture was then separated into 5 ml aliquots in 15 ml tubes and centrifuged at 2E+03 rpm for 5 minutes to obtain biofilm pellets. The supernatant was decanted and the pellets were resuspended 1 ml of deionized water to wash the biofilm to eliminate salts from the media. The resuspended biofilm was centrifuged for a second time at 2E+03 rpm for 5 minutes. The 1 ml biofilm pellet was then transferred to an upper insert of a transwell containing the desired cobalt concentration in the lower part of the well. The material was left at the desired temperature and pH for the set exposure time before the upper transwell insert was removed and the remaining cobalt concentration in the free solution was measured. 12-well transwell plates with a polycarbonate 0.45 *µ*m membrane size (VWR) were used for all tests. For pH experiments, sodium dihydrogen phosphate anhydrous / phosphoric acid buffers at pH values of 2 and 9 were used to tune the pH to the desired values. Following metal exposure, 50 *µ*l of biofilm from the upper well was used to make serial dilutions and was bead plated on LB agar plates to measure the viability of cells following metal exposure.

### Cobalt Measurement

The cobalt concentration remaining in the free solution following biofilm exposure was determined using a colorimetric reaction. A sample volume of 600 *µ*l was added to a 50 ml beaker along with 3 ml of a 40% weight/volume sodium citrate in water solution. An indicator solution made up of 0.1% bromothymol blue in 50% ethanol was prepared in advance. A volume of 0.3 ml of indicator solution was then added to the reaction. The solution was neutralized with 10% sodium hydroxide solution added dropwise until the color matched a standard solution (20 ml water, 5 ml 40% sodium citrate solution, and 0.5 ml indicator solution). A volume of 6 ml of 0.5% w/v nitroso-R-salt in deionized water was then added. The solution was brought to a boil and 3 ml of 34% nitric acid was added. Following 1 minute of boiling, the solution was cooled to room temperature and a 200 *µ*l aliquot was transferred to a 96-well plate for measurement on a plate reader at 500 nm. The absorbance of a sample blank without cobalt was subtracted from the sample absorbance value before comparing the sample absorbance against cobalt standard solutions to determine the concentration of cobalt.

### Zeta Potential and Molecular Weight Measurements

Zeta potential and *M_W_* were measured using a Malvern Nano Zs Zetasizer (Malvern Pananalytical, Malvern, U.K.) following the protocol used previously.[65] Solutions were given 2 hours to equilibrate before measurement in a polystyrene or zeta potential folded capillary cuvette for measurement of either *M_W_* or zeta potential respectively. Zeta potential was measured at a concentration of 0.1% with deionized water as the buffer. The *M_W_* was determined using the light scattering measurements from five concentrations, 0%, 0.025%, 0.05%, 0.075%, and 0.1%. The Rayleigh equation was then applied to calculate the *M_W_* of the sample.[39]

### Fourier-Transform Infrared Spectroscopy

Fourier-Transform Infrared Spectroscopy (FTIR) measurements were conducted using a Bruker Vertex FT-IR spectrometer (Bruker Corp., Billerica, MA). A spectral range of 600 cm^-1^ to 3500 cm^-1^ was selected. Approximately 5 mg of sample were loaded into the sample compartment to completely cover the attenuated total reflectance (ATR) crystal. A nitrogen atmosophere was used for all reference and sample measurements.

### Rheometry

Rheometry testing was done using a TA Instruments DHR3 rheometer (Texas Instruments, Dallas, TA). A strain sweep at a frequency of 1 rad s^-1^ was used to establish the range of the the linear viscoelastic region. A strain of 1% was selected to be used for frequency sweeps based on the results of the strain sweeps. A sweep of strain and frequency was completed for every sample. A 20 mm parallel plate fixture was used with the gap set to 300 µm, requiring a sample volume of approximately 100 µl. Biofilm samples were exposed to desired temperature or pH conditions for 24 hours in a transwell plate before being harvested for rheology. The biofilm concentration was 2 mg / ml of biofilm dry weight to ml of water. The parallel plates were thoroughly cleaned before use and between samples with deionized water and isopropyl alcohol and dried with compressed air. The sample was added onto the lower plate and the top fixture lowered to the set gap value. Graphs presented are the average of three independent trials.

### Alcian Blue Staining

Collection of floating biofilms: Overnight liquid cultures were diluted 100-fold into 250 mL glass beakers containing 200 mL of PYE medium. Cultures were grown statically at room temperature for 5 days to allow for biofilm maturation. Four layers of cheesecloth were secured over the opening a 1 L glass beaker with a rubber band, and biofilm cultures were carefully poured over the cheesecloth to collect the biofilm matrix. The gelatinous filtrate was then collected with a metal spatula and transferred into a 15 mL conical tube. SDS-PAGE analysis of biofilm composition: 4x Laemmli buffer (250 mM Tris-HCl pH 8.0, 8 % w/v SDS, 40 % v/v glycerol, 0.05 % w/v Bromophenol Blue and 5 % v/v betamercaptoethanol) was added to biofilm extracts to achieve a final concentration of 1x. Solubilized extracts were serially diluted, heated at 95*^◦^* for 5 min and loaded onto 4-20 % polyacrylamide gradient gels (BioRad). After resolving, gels were fixed in a solution of acetic acid/ethanol/water (1:5:4) for 1h and subsequently washed for 1 h in deionized water. The fixing and washing steps were repeated a second time. Gels were then stained overnight in a 0.2 % solution of Alcian Blue, 3 % v/v acetic acid and 50 mM MgCl_2_. A solution of 3 % v/v acetic acid and 50 mM MgCl_2_ was then used to destain the gels.

### Cobalt Cell Uptake Test

Cells were separated from the secreted biofilm to compare the amount of cobalt removed from solution by the biofilm alone and by the cells. Centrifugation of 5 ml of culture at a maximum possible speed of 7830 rpm (Eppendorf, Germany) for 15 minutes was used to separate out cells from the biofilm. A volume of 1 ml of biofilm was then carefully pipetted off and placed in a separate transwell plate for cobalt exposure. The 100 µl cell pellet was resuspended in 0.9 ml of water and added to a transwell plate well for cobalt exposure. Both the biofilm and the cells underwent 24 hours of exposure to water with a cobalt concentration of 2E+03 µg/L. The concentration of cobalt remaining in the water was then measured following the exposure. One group of exposed cells were also exposed to 5 mg/ml for 6 hrs to lyse cells. The cobalt concentration in the free solution was measured after the lysozyme exposure. Cells were found to survive lysozyme exposure if not first separated from the biofilm.

### Cobalt Recovery Tests via Degrading Protein

Cobalt recovery was conducted using guanidinium iodide and Proteinase K to breakdown metal binding proteins. A concentration of 10 % guanidinium iodide (MS140000, Greatcell Solar, Australia) Materials,) and 0.2 % Proteinase K (20E0456058, VWR, Radnor, PA) respectively for 6 hours. The metal uptake with and without the chemical treatments were then measured to compare the effect of degrading proteins on metal uptake and recovery.

### Column Uptake Test

Biofilm pellets with a dry weight of 160 mg were produced by centrifuging culture at 2E+03 rpm for five minutes. The pellets were washed with deionized water before being transferred to a coffee filter that had been pre-wetted with deionized water. Four volumes of cobalt contaminated water at a concentration of 2E+03 *µ*g/L were run through the filter with the pellet. Filtration was gravity driven and each 10 ml volume was collected in a unique 50 ml Falcon tube placed under the filter. The cobalt concentration of each 10 ml volume was then measured. Tests were performed in triplicate. Results were compared to a sample of deionized water run through the biofilm filter as well as a cobalt sample run through just the coffee filter to eliminate variation in absorbance that were due to cell debris or cobalt adsorption of the coffee filter.

### Batch Uptake Test

Liter scale uptake of cobalt was demonstrated by exposing cobalt contaminated water to biofilm for 24 hours. Tests were conducted in liter sized glass bottles. The concentration of cobalt tested was 2E+03 *µ*g/L with 0.2 mg of biofilm per 1 ml of water (0.2 g total biofilm used per each liter container). A cobalt stock solution was made from cobalt (II) chloride salt at a concentration of 40 g/L. The stock solution was then diluted to make liter volumes of cobalt After exposure, a 50 ml water sample was taken and filtered to remove any biofilm or cell debris before measuring cobalt uptake from the free solution. Proteinase K was then added for a 6 hour exposure at a concentration of 2 mg/L to digest proteins involved in metal uptake and recover the cobalt. Following the Proteinase K exposure, another 50 ml water sample was extracted, filtered, and used for cobalt measurement to calculate metal recovery. All tests were performed in triplicate.

## Acknowledgement

This work was supported through the NSERC CGS-D scholarship (E. W. v. W.), ELI grant (M. P. B), Department of Energy (Grant DE-SC0025354) (M. N. S. and E. W. v. W.), the Beckman Young Investigator award (D. M. H.), and NIH R35GM150652 (D. M. H). This work was supported and performed in part at the Engineered Living Materials institute, Cornell Center for Materials Research shared instrumentation facility, and the Cornell NanoScale Facility, a member of the National Nanotechnology Coordinated Infrastructure (NNCI), which is supported by the National Science Foundation (Grant NNCI69 2025233).

## Supporting Information Available

Supporting Information: Additional experimental details and photos including growth curves, rheological strain sweeps, FTIR-ATR spectra of the biofilm, and all trials for metal removal and viability.

