## Supplementary Figures for "Rheinheimera sp. T2C2 Bacterial Biofilm for Bioremediation of Cobalt (II)"

### Supporting Figures

#### 1.0 Ideal Characteristics of *Rheinheimera sp.* T2C2 for Heavy Metal

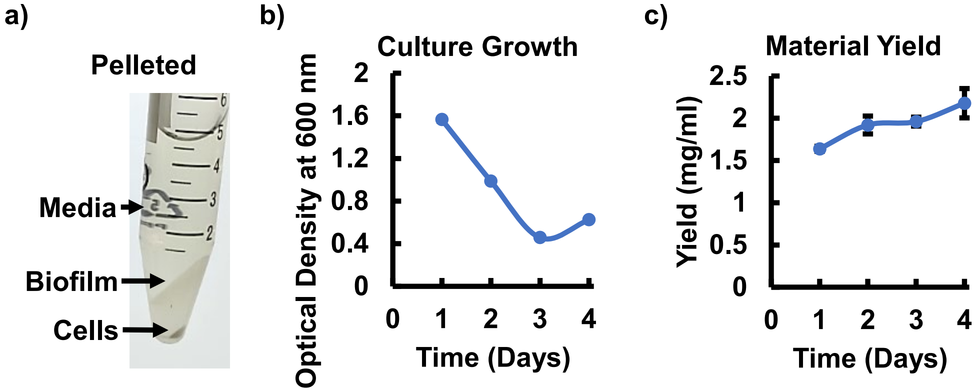

Figure S1: a) The biofilm pellet can be grown and collected via centrifugation of the culture. b) Optical density and c) yield of *Rheinheimera sp.* T2C2 over a four day growth period before biofilm is harvested.

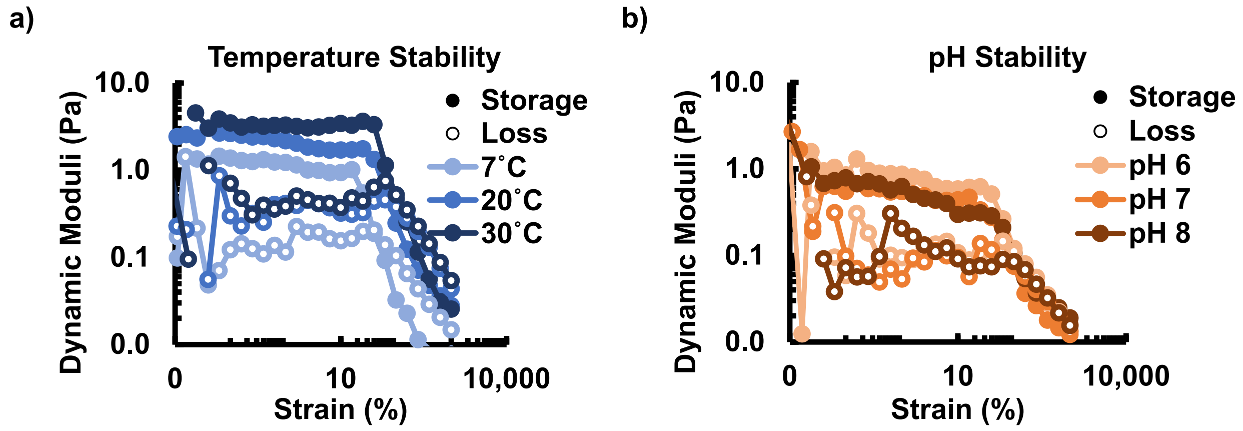

Figure S2: Rheological strain sweeps for a) temperature and b) pH water condition tests conducted at a frequency of 1 rad/s.

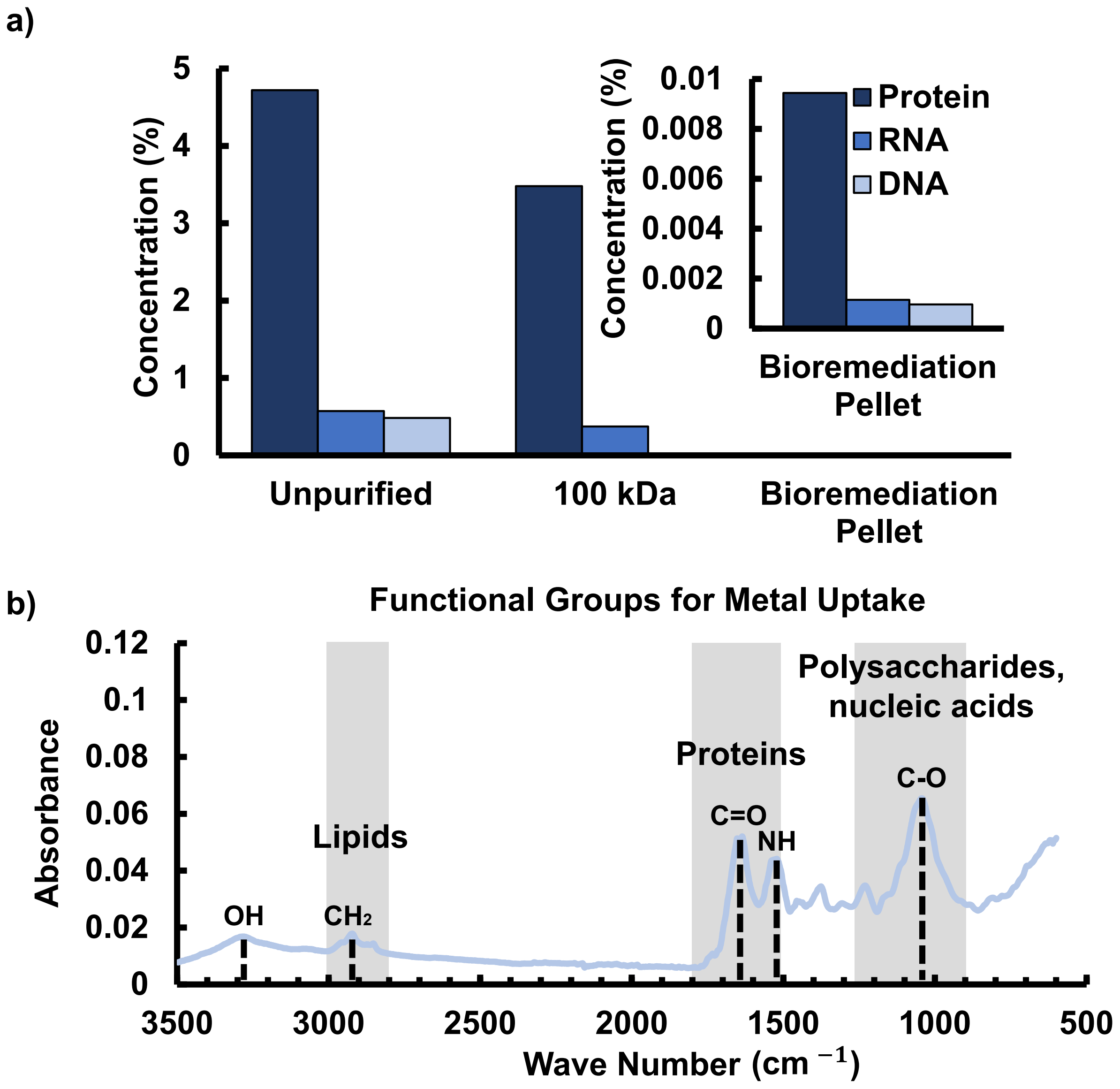

Figure S3: a) Protein, RNA, and DNA composition of *Rheinheimera sp.* T2C2 biofilm measured using a Qubit fluorometer. Inset shows the values for the bioremediation pellet to facilitate visualization. Dialysis results in a loss of low molecular weight small molecules and proteins. b) The biofilm consists of polysaccharides, proteins, and lipids with functional groups that may be suited for heavy metal binding as identified using FTIR-ATR.

#### 2.0 Biosorption of Cobalt using *Rheinheimera sp.* T2C2

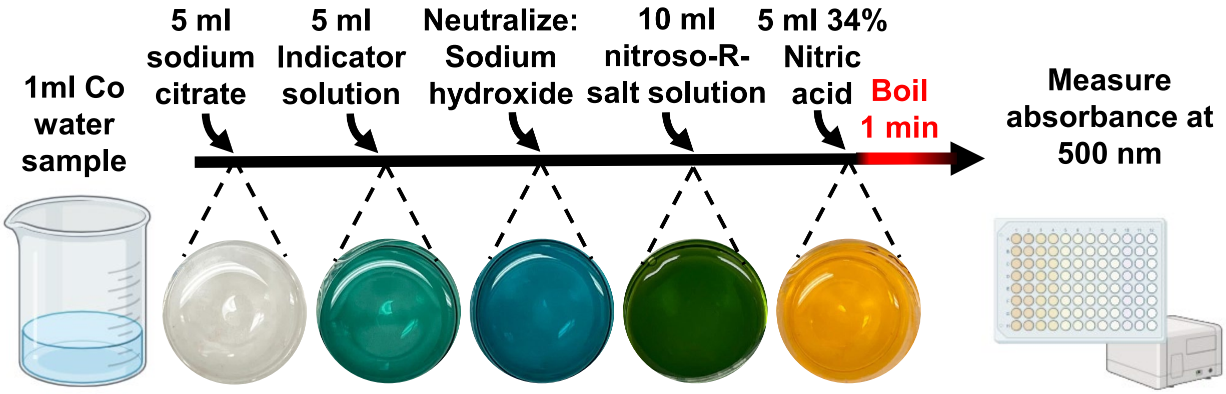

Figure S4: Colorimetric cobalt measurement assay using nitroso-R salt.

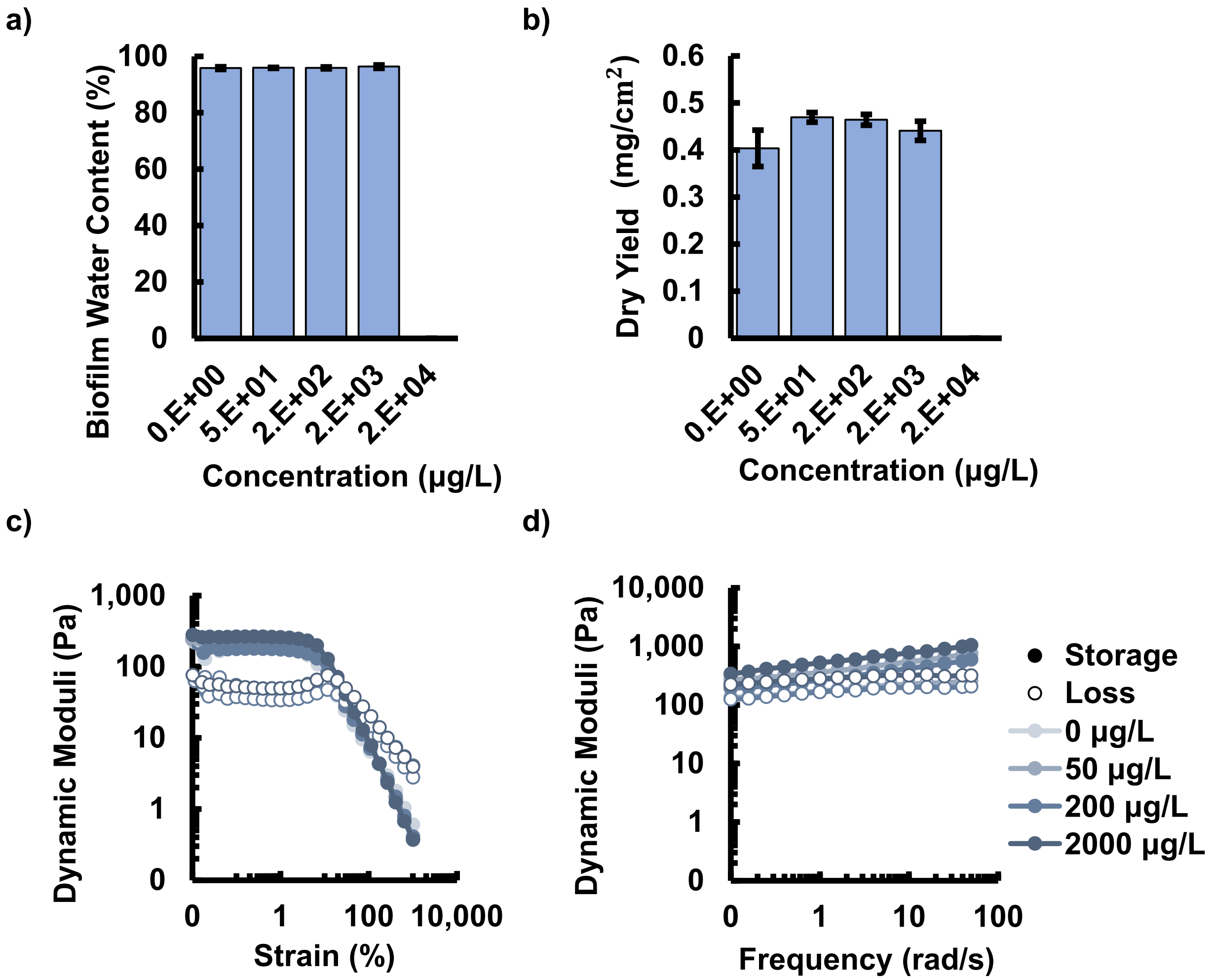

Figure S5: The effect of cobalt concentration for *Rheinheimera sp.* T2C2 biofilm grown on cobalt-LB-agar plates. Cobalt concentrations below 20000 $\mu$g/ml had minimal effect on a) biofilm water content, b) yield and rheological properties including c) strain sweep at 1 rad/s and d) frequency sweep at 5% strain.

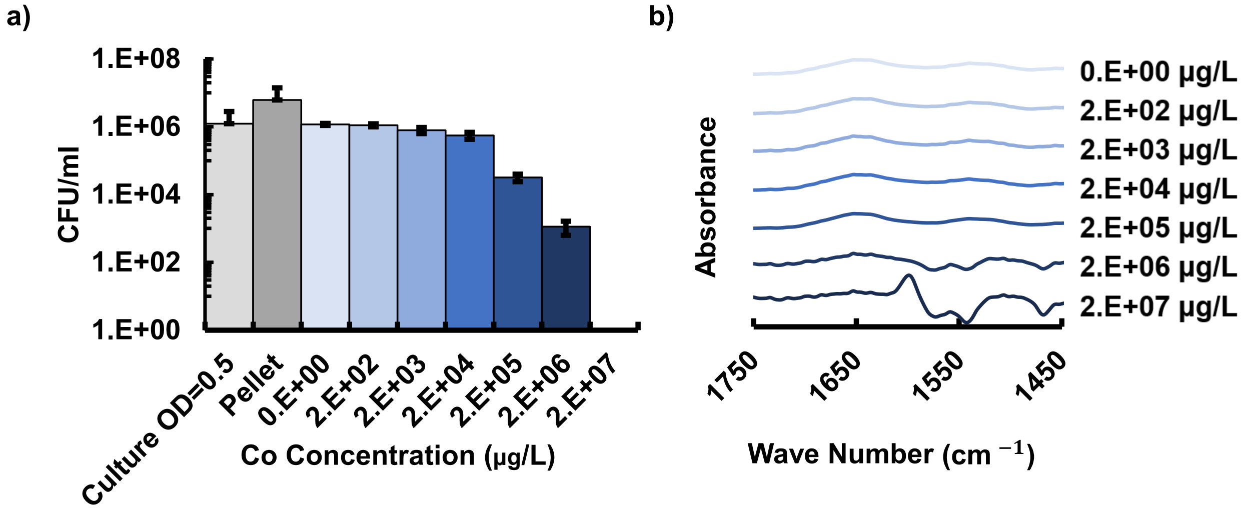

Figure S6: a) Cell viability decreased with increasing metal concentration. b) Changes in ATR-FTIR spectra were observed for increasing concentration of cobalt. All error bars report standard deviation (n=3).

#### 3.0 The Effect of Varied Water Conditions on Biofilm Biosorption of Cobalt

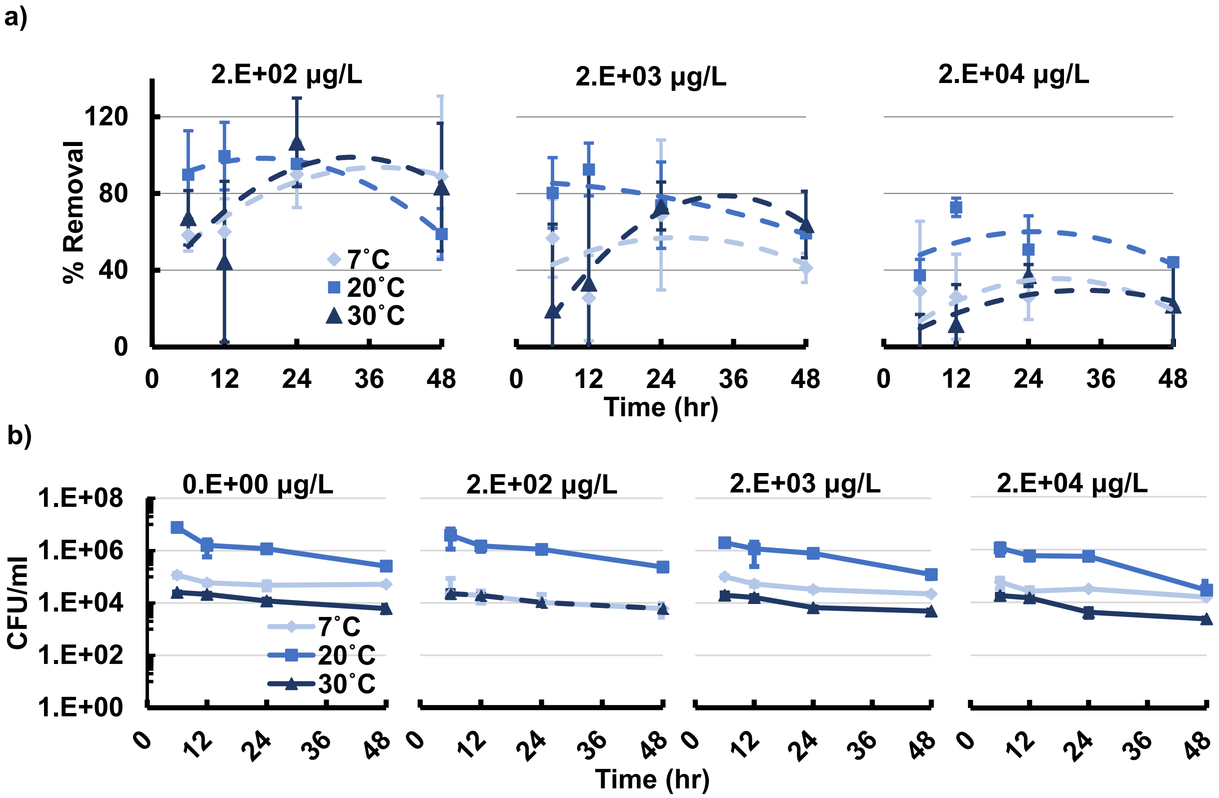

Figure S7: a) The effect of varying water temperature and exposure time on percent cobalt removal showed differing peak uptake times depending on temperature. The percentage metal removed decreased for increasing metal concentration. b) Cell viability in varied water temperatures indicated that T2C2 grows best at a temperature of 20°C. Viability decreases with increased metal exposure time for all temperatures tested. Error bars show standard deviations (n=3).

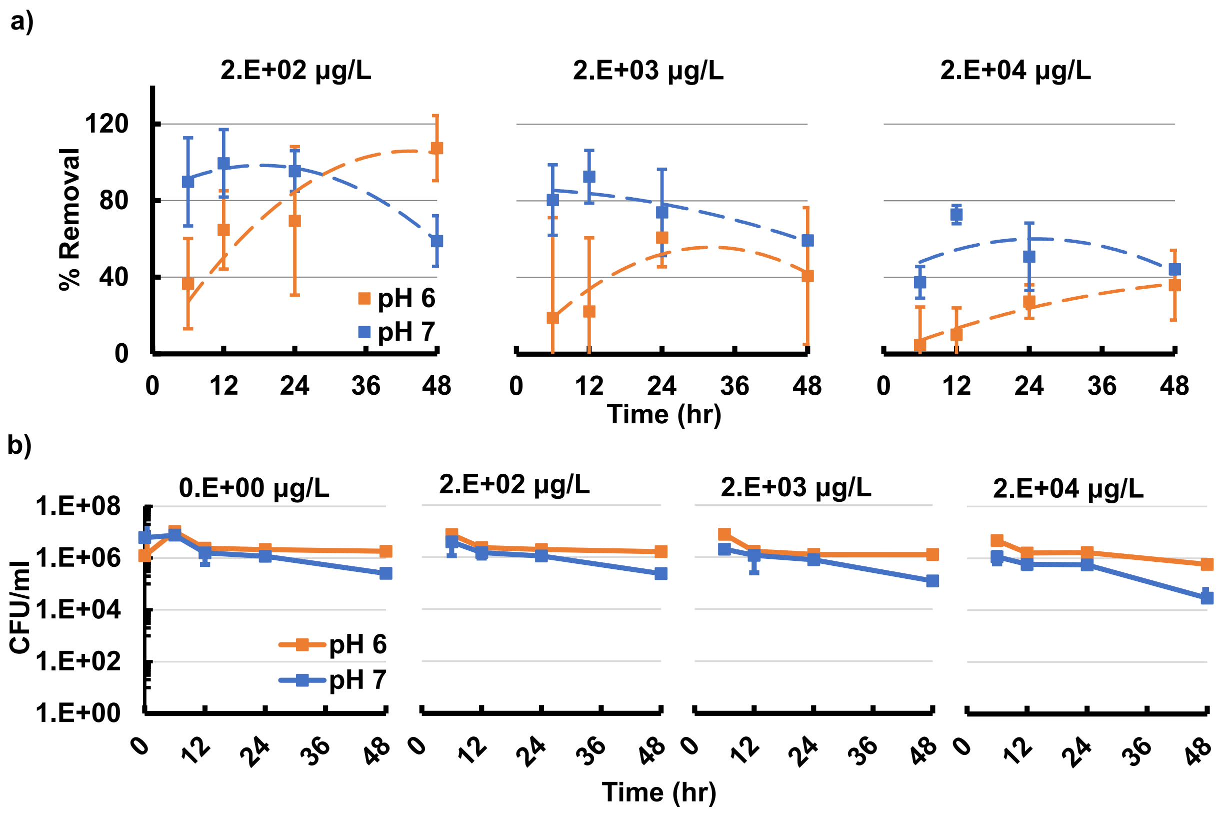

Figure S8: a) Varying water pH and exposure time altered the percent cobalt removal producing different peak uptake times. The percentage metal removed decreased for increasing metal concentration. A higher pH of 7 generally resulted in higher metal uptake that a pH of 6. b) Cell viability for varied water pH indicated similar growth with decreased viability for longer metal exposure times. Error bars show standard deviations (n=3).

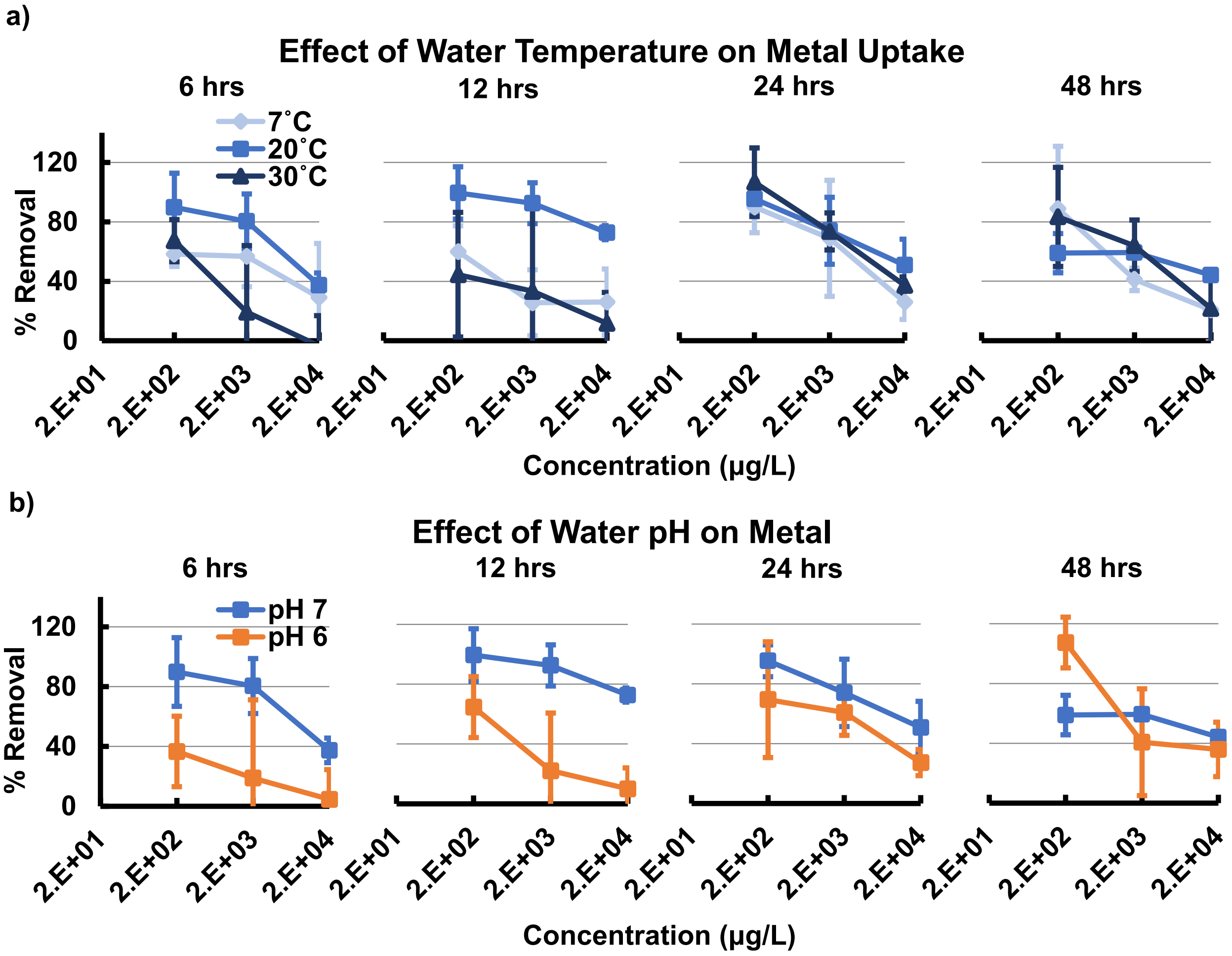

Figure S9: Percent of cobalt removal vs, initial concentration for 6, 12, 24, and 48 hours for a) varied water temperature and b) pH. Error bars show standard deviations (n=3).

#### 4.0 Biofilm Mode of Metal Uptake

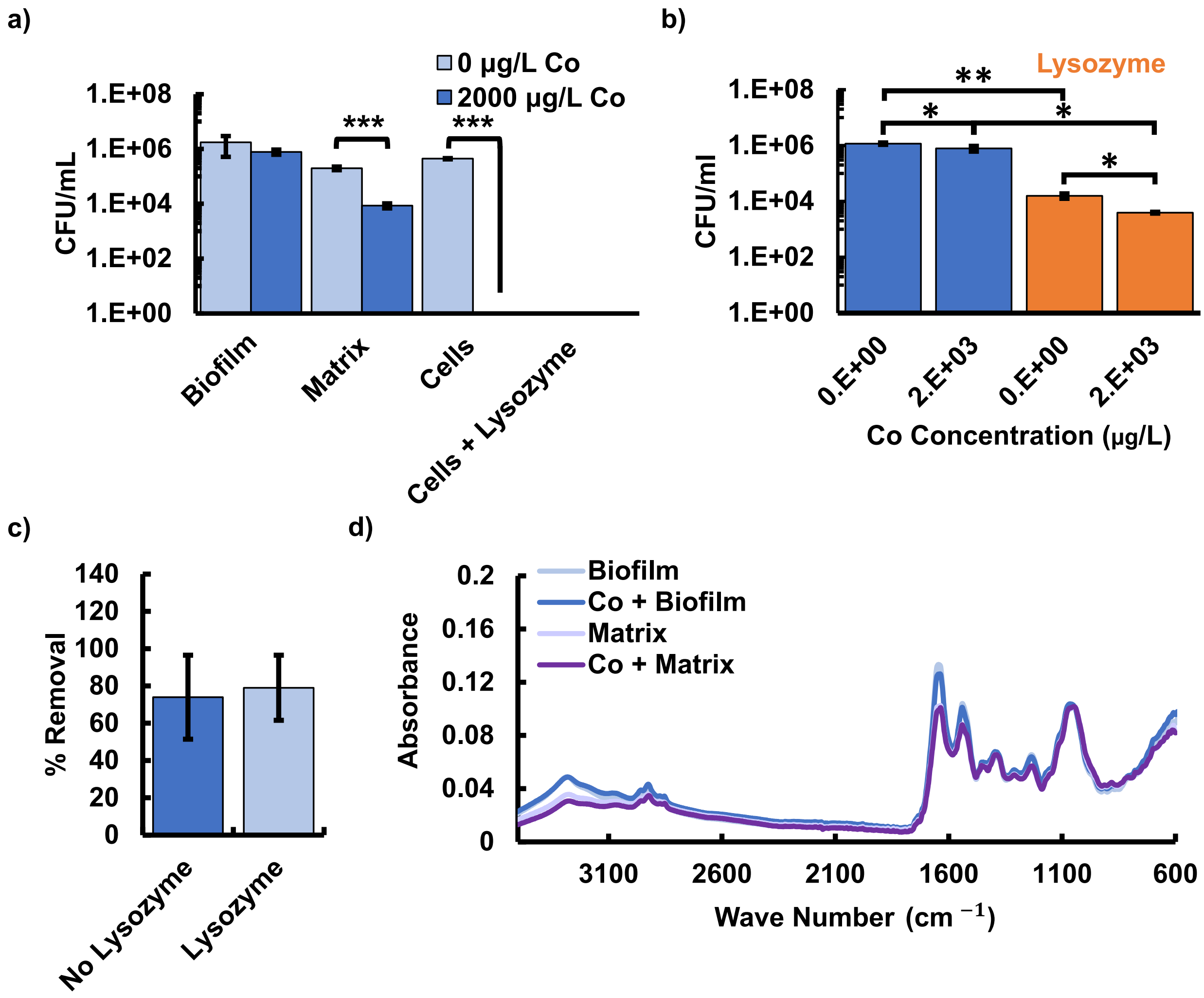

Figure S10: a) Cell viability tests for centrifugation to separate the cells from the biofilm. Note that no cells survived lyophilization of the biofilm. b) Cell viability due to a 6 hr lysozyme treatment, without first separating cells, at a concentration of 5 mg/ml resulted in the majority of cells surviving. c) No statistically significant change in metal removal was observed when cells were treated with lysozyme without initial cell-film separation. All error bars show standard deviations (n=3). d) FTIR results for biofilm with cells and matrix without cells exposed to cobalt concentrations of 0 and 2000 $\mu$g/L, normalized to carbohydrate peak at 1045 cm$^{-1}$.

#### 5.0 Cobalt Recovery via Biofilm Protein Degradation

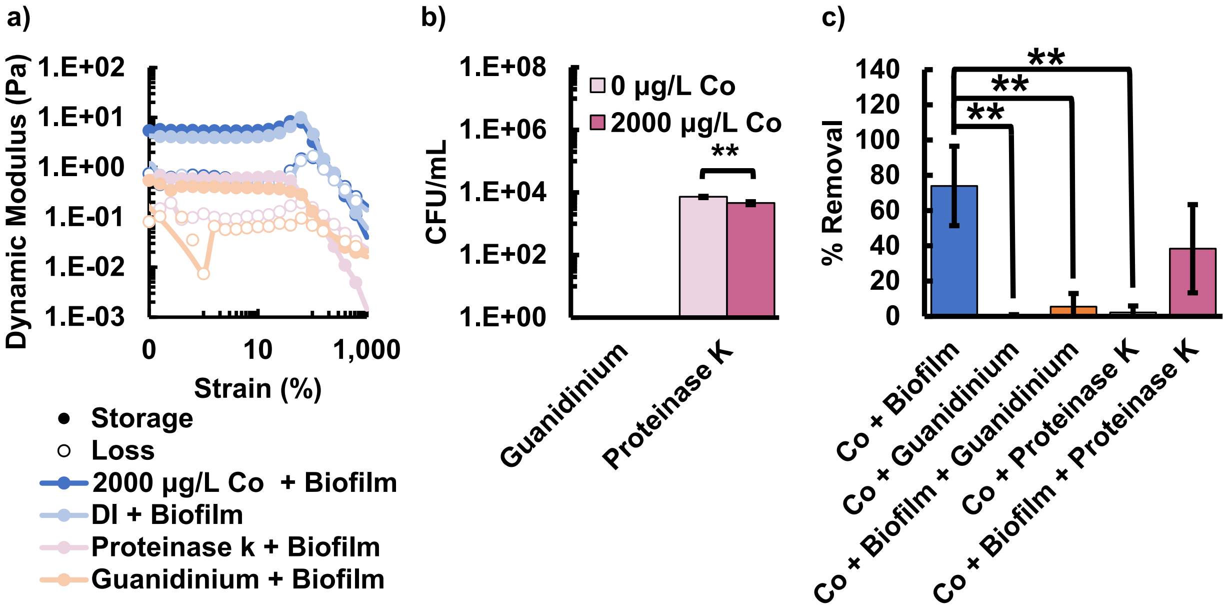

Figure S11: Metal recovery from the biofilm was conducted via protein degradation a) Strain sweep showing biofilm degradation due to proteinase k and guanidinium exposure. b) Cell viability for biofilm degradation using proteinase K (0.2% w/v) and guanidinium iodide (10% w/v) for 6 hours respectively. c) percent removal for biofilm and treated groups demonstrating uptake and metal release into free solution. Error bars show standard deviations (n=3).
